# Functional linkage of gene fusions to cancer cell fitness assessed by pharmacological and CRISPR/Cas9 screening

**DOI:** 10.1101/559690

**Authors:** Gabriele Picco, Elisabeth D Chen, Luz Garcia Alonso, Fiona M Behan, Emanuel Gonçalves, Graham Bignell, Angela Matchan, Beiyuan Fu, Ruby Banerjee, Elizabeth Anderson, Adam Butler, Cyril H Benes, Ultan McDermott, David Dow, Francesco Iorio, Euan Stronach, Fengtang Yang, Kosuke Yusa, Julio Saez-Rodriguez, Mathew J Garnett

## Abstract

Many gene fusions have been reported in tumours and for most their role remains unknown. As fusions can be used clinically for diagnostic and prognostic purposes, and are targets for treatment, it is crucial to assess their functional implications in cancer. To investigate the role of fusions in tumor cell fitness, we developed a systematic analysis utilising RNA-sequencing data from 1,011 human cancer cell lines to functionally link 8,354 gene fusion events with genomic data, sensitivity to >350 anti-cancer drugs and CRISPR-Cas9 loss-of-fitness information. Established clinically-relevant fusions were readily identified. Overall, functional fusions were rare, including those involving cancer driver genes, suggesting that many fusions are dispensable for tumor cell fitness. Novel therapeutically actionable fusions involving *RAF1, BRD4* and *ROS1* were verified in new histologies. In addition, recurrent *YAP1-MAML2* fusions were identified as activators of Hippo-pathway signaling in multiple cancer types, supporting therapeutic targeting of Hippo signalling. Our approach discriminates functional fusions, identifying new drivers of carcinogenesis and fusions that could have important clinical implications.

**Significance:** We identify fusions as new potential candidates for drug repurposing and drivers of carcinogenesis. These results support histology agnostic marker-driven precision cancer medicine. Most fusions are not functional with implications for interpreting cancer fusions reported from clinical sequencing studies.

## Introduction

Oncogenic gene fusions occur in solid tumours and hematologic malignancies, and are used for diagnostic purposes, patient risk stratification and for monitoring of residual disease^1^. Critically, the chimeric protein encoded by fusions may be a tumor specific target for treatment, resulting in significant clinical benefit for patients^2,3^. Fusions are often associated with a specific tissue histology, but can occur at a low frequency in multiple histologies. Gene fusion transcripts are composed of two independent genes formed either through structural rearrangements, transcriptional read-through of adjacent genes, or pre-mRNA splicing. The exchange of coding or regulatory sequences between genes can result in aberrant functionality of the fusion protein, and deregulation of the partner genes, including overexpression of oncogenes and decreased expression of tumor suppressor genes (TSG).

Discriminating between fusions that have a role in cancer fitness and those that do not is a major challenge with important clinical implications^4^. Deep sequencing technology together with sensitive fusion detection algorithms have led to a dramatic increase in the number of reported cancer-associated fusions^5^. Most fusion transcripts are likely the indirect consequence of genomic instability or false-positive events due to error-prone fusion calling. Previous studies have focused on the identification of fusions, or have investigated the function of specific gene fusions, for example in the setting of acute myeloid leukemia (AML)^6^. The functional role of most fusions has not been investigated.

We have generated large-scale genomic and pharmacological datasets for >1,000 human cancer cell lines as part of the Genomics of Drug Sensitivity in Cancer (GDSC) project^7,8^. These datasets, together with recent advances in CRISPR genetic screening technology, make it now possible to systematically assess the contribution of fusions transcripts to cancer cell fitness.

Here, we report the first comprehensive functional landscape of fusions events using RNA sequencing (RNA-seq) data for 1,011 human cancer cell lines. We investigate the functional relevance of gene fusions using differential gene expression, drug sensitivity to >350 anti-cancer compounds, and whole-genome CRISPR-Cas9 drop out screens to identify fusions required for cancer cell fitness. To our knowledge, this study is the first large-scale systematic analysis in a large collection of human cancer models to unveil the largely unexplored functional role of gene fusion.

## Results

### Landscape of fusion transcripts

To systematically identify gene fusions in diverse cancer types, we first analyzed RNA-seq data to define fusion transcripts in the GDSC cancer cell lines (1,034 samples from 1,011 unique cell lines) representing 41 cancer types (Fig. 1a). RNA-seq data for 587 cell lines was obtained from the Cancer Genome Hub (CGHub) and 447 cell lines were sequenced at the Sanger Institute. Fusion calling algorithms are prone to detecting false positives from sequencing artefacts and alignment ambiguities^9^. To improve the accuracy of fusion transcript calling, we used three different algorithms, deFuse, TopHat-Fusion and STAR-Fusion, across all samples^10–12^ and applied stringent filtering criteria. In total, 10,514 fusion transcripts were called by more than one algorithm and taken forward for this study (Fig. 1b). Targeted PCR of 406 putative fusion breakpoints resulted in validation rate of 71.6%. Furthermore, we compared the 23 samples with RNA-seq data from both Sanger Institute and CGHub (Supplementary Table 1) and the proportion of fusions transcripts in both data sources for a given cell line was 70.4%.

**Figure 1:**
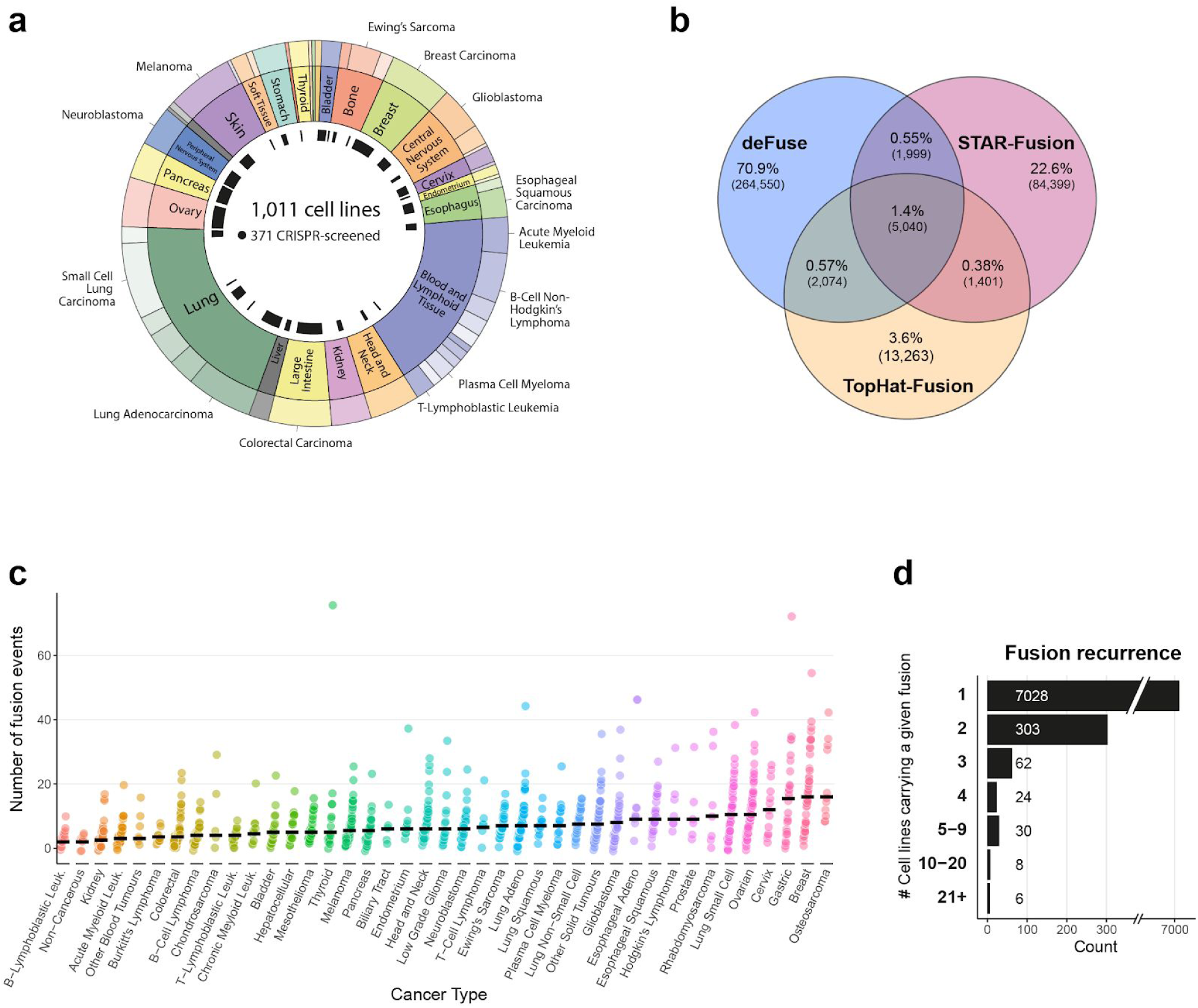
Landscape of gene fusions in cancer cell lines. (a) Summary of cancer types represented by the cell lines and CRISPR dataset used for this study. (b) Venn diagram showing overlap in fusion transcript calls using three algorithms, DeFuse, STAR-Fusion and TopHat-Fusion. (c) Frequency of gene fusions events identified in cancer cell lines, separated by cancer type. (d) Fusion event recurrence in the cancer cell lines.

Some fusions have multiple transcripts in the same cell line and so we define a ‘fusion event’ as a fusion present in a cell line. Thus, we identified 10,514 fusion transcripts, representing 8,354 gene fusion events and, because a small number of fusions are recurrent, 7,430 unique fusions (Supplementary Table 2).

Next, we examined the number of fusion events that occurred in different cancer types. Cell lines had a median of six fusion events and 26% of fusion events were predicted to be in-frame. Fusion numbers varied by cancer type (Fig. 1c), with osteosarcoma and breast cancer having the most (median of 16 fusion events per cell line), and kidney cancers and B-lymphoblastic leukemia together with three non-cancerous immortalized human cell lines having the lowest median number of fusion events (median = 2). The prevalence of fusion events for each cancer type in our cell lines was slightly higher, but significantly correlated with the frequency reported from the analysis of 9,624 patient samples (p < 0.001, R^2^ = 0.42; Supplementary Fig. 1), indicating cell lines reflect the frequency of fusions in tumours from different tissues^13^. We identified recurrent known oncogenic fusions events in our dataset, including *BCR-ABL1* (n = 11), *NPM1-ALK* (n = 5), *EWSR1-FLI1* (n = 24) and *TMPRSS2-ERG* (n = 2). Of note, only 431 of 7,430 (6%) fusions were recurrent, while the remaining were detected in only one cell line (Fig. 1d), indicating most fusions are rare.

Of the fusion events we identified, 11.1% have been reported previously in human tumor samples^13^ and for 14.2% of the fusion events, at least one of the fused genes was found in the COSMIC Cancer Gene Census, representing a significant enrichment for cancer genes (odds ratio = 1.8 and p < 0.001, Fisher’s test). TSGs were enriched as 5’ end partner genes (odds ratio = 2.1 and p < 0.001, Fisher’s test), while oncogenes were enriched as 3’ or 5’ genes (odds ratio = 2 and 1.8, respectively, p < 0.001, Fisher’s test). We found known oncogenic fusions enriched in specific cancer types consistent with their pathognomonic nature, such as *ABL1*-fusions in chronic myeloid leukemia (n = 9; p = < 0.001), *EWSR1-FLI1* fusions in Ewing’s sarcoma (n = 24; p = < 0.001) and *FGFR3* fusions in bladder cancer (n = 3; p = < 0.001) (Supplementary Fig. 1). In summary, using stringent criteria we built a comprehensive landscape of fusions in cancer cell lines, most of which occur at a low frequency, and reflect the prevalence and tissue specificity in tumor samples.

### Fusion transcripts impact gene expression

Fusions may result in altered expression of either or both of the fusion partner genes^13^. To identify genes whose expression is altered when fused, we first aggregated fusion events that had a common gene partner at the 5’ or the 3’ end to increase sample size and statistical power. We then used a linear regression model to link expression with the presence of a fused gene, incorporating bias due to copy number alterations and cancer type. In total, we tested 902 genes (5’ genes: 611 and 3’ genes: 383) that involved 3,048 fusions. We identified 172 (19%) genes significantly associated with differential expression (5’ genes: 54 (9%) and 3’ genes: 118 (31%)) that encompassed 592 fusions. Of the significantly associated genes, 24 (14%) were known cancer drivers from the COSMIC census (2.5% of the total; Fig. 2a and Supplementary Table 3). As expected, several TSG such as *TP53, APC* and *KDM6A* were significantly associated with reduced expression (p < 0.001, Supplementary Fig. 1). In contrast, many known oncogenes fused at the 3’ were overexpressed, including *ALK, ERG, FL1, MYC, MLL4* and *ROS1* (p < 0.001, Supplementary Fig. 1).

**Figure 2:**
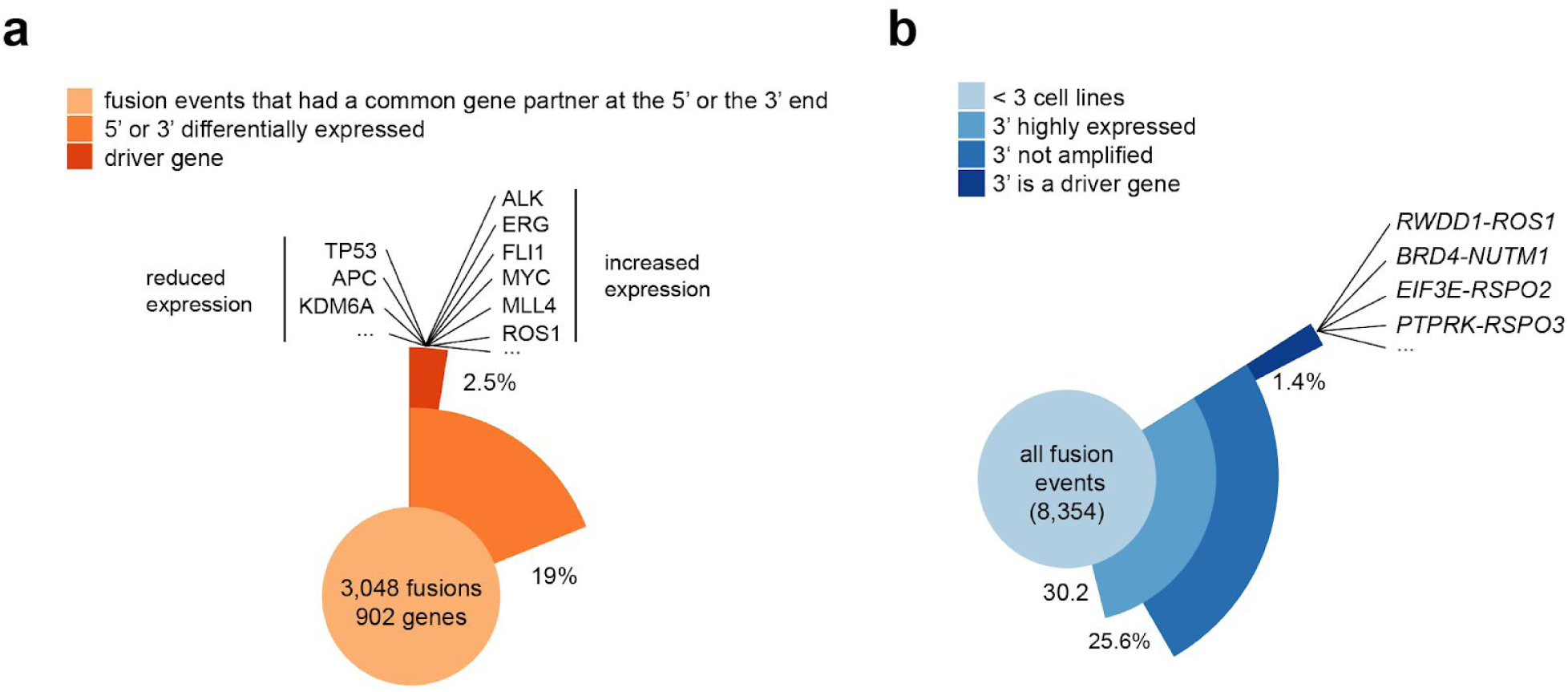
Fusion transcripts impact gene expression. (a) Frequency of a statistical association between the presence of a recurrent fusion (n > 2 cell lines) and differential gene expression. Examples of downregulated TSG and overexpressed oncogenes are displayed. (b) Frequency of co-occurrence of a gene fusion and overexpression of the 3’ fusion gene for each fusion event. RSPO2, RSPO3 and NUTM1 are examples of overexpressed cancer driver genes involved in previously unreported gene fusions.

Because most fusions are rare and therefore not suitable for linear regression modelling, we also annotated expression of genes involved in each fusion event (n = 8,354). We focused on 3’ end genes with exceptionally high expression because overexpression of proto-oncogenes occuring as 3’ partner genes is observed in several malignancies^13,14^. We found that 25.6% (n = 2,145) of fusion events were coincident with high expression and did not co-occur with copy number amplification. Only 5.4% (1.4% of the total; n = 117) of these fusion events involve the overexpression of a driver gene (Fig. 2b) (Supplementary Table 4). This analysis revealed that aberrant transcript expression of genes involved in gene fusions is a common event, but only a small subset of these fusions involve established driver oncogenes.

Novel contexts of chimeric transcripts were identified leading to overexpression of known cancer genes located the fusion 3’ end, such as *NUTM1, RSPO2/3* and *ROS1* (Fig. 2b). In support of this observation, we validated by Sanger sequencing and fluorescent in situ hybridisation (FISH) a previously uncharacterized *RWDD1-ROS1* fusion in the OCUB-M cell line, which is derived from a triple-negative breast cancer (Supplementary Fig. 2). ROS1 is a receptor tyrosine kinase and gene rearrangements leading to ROS1 overexpression have been identified and validated as therapeutic biomarker of response to ROS1 kinase inhibitors in non-small cell lung cancer and, recently, in other cancer types (Supplementary Fig. 2)^15^. The fusion retains the ROS1 protein kinase domain and OCUB-M cells display sensitivity to crizotinib and foretinib, two potent ROS1 inhibitors (Supplementary Fig. 2)^16,17^. Interestingly, in a dataset of 590 breast cancer patients, we identified a triple-negative and a HER2+ sample carrying in-frame fusions involving the ROS1 kinase domain^18^ (Supplementary Fig. 2), suggesting that a rare subset of breast cancer patient could be potentially eligible to targeted tyrosine kinase inhibitor-based therapies.

### Systematic analysis for fusion markers of drug response

Fusion proteins can impact on clinical responses to therapy. Consequently, we reasoned that differential drug sensitivity in cell lines could be used to identify functional fusions, as well as opportunities for repurposing of existing drugs. We used an established statistical model^8,19^ to perform an analysis of variance (ANOVA) linking the 431 recurrent gene fusions (n ≥ 2 cell lines) with 308,634 IC50 values for 409 anti-cancer drugs (334 unique compounds) screened across 982 of our cell lines as part GDSC project (Fig. 3a, Supplementary Table 5). The compounds assessed consisted of anti-cancer chemotherapeutics and molecularly targeted agents, including many which are FDA-approved (n = 46; Fig. 3a) or in clinical development (n = 65). This included data for 155 new compounds and a total of 212,774 previously unpublished IC50 values. Preliminary analyses indicated that mutations in cancer driver genes co-occurring with fusions in cell lines were frequent confounders when identifying fusion-specific associations. To control for this, we first identified associations between 717 cancer driver mutations and copy number alterations and drug sensitivity, then used them as a covariate in the ANOVA to identify fusion-specific associations (Supplementary Table 6). Adding the covariates resulted in 11 fusion associations falling below our threshold for statistical significance. For instance, the association of *NKD1-ADCY7* with BRAF-inhibitor dabrafenib was explained by the presence of a *BRAF* mutation in one highly sensitive cell line (Supplementary Fig 3).

**Figure 3:**
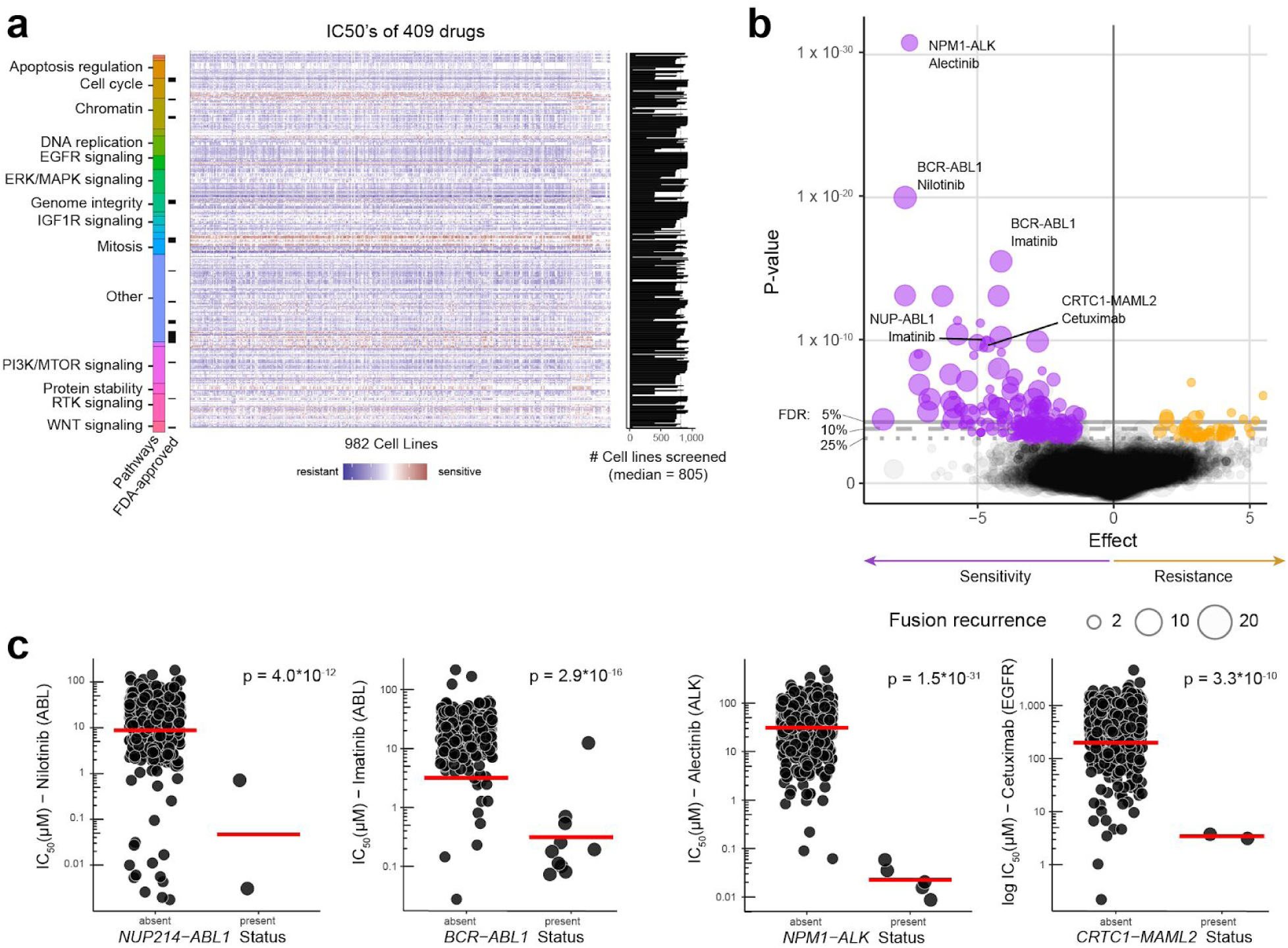
Gene fusions as therapeutic biomarkers. (a) Overview of GDSC drug sensitivity data (reported as IC50 values), including number of cell lines screened per drug (median = 805) and FDA-approval status of compounds. Compounds (n = 409) are grouped by target or pathway. (b) Volcano plot of ANOVA result (y-axis: p-value) for significant fusion–drug associations. Each circle represents an association, with circle size the number of cell lines harbouring the associated fusion event (fusion recurrence). Negative effect sizes are associated with sensitivity and positive effect sizes resistance. Representative fusions-drug associations are labelled. (c) Examples of differential drug sensitivity in cell lines stratified by fusion status. Nominal therapeutic drug targets are in brackets. Each circle is the IC50 for an individual cell line and the red line is the geometric mean. Association significance (p-values) shown are from the ANOVA test.

We identified 227 large-effect size associations (FDR < 25% and Glass Deltas > 1; the Glass Delta is a measure of effect size incorporating the standard deviation) between gene fusions and drug sensitivity (Fig. 3b; Supplementary Table 7). At the level of individual fusion events, 284 (21%) of 1,355 tested fusion events showed a significant association with a drug. Most of the the strongest fusion-drug associations were well understood cases, such as sensitivity of *ALK*-fusion positive cell lines to ALK inhibitors, for example, alectinib, (FDR < 0.1%), and sensitivity of *BCR-ABL1* translocation positive cells to ABL inhibitors, such as imatinib and nilotinib, (FDR < 0.1%) (Fig. 3c). We also identified associations with low frequency fusions, such as sensitivity to multiple EGFR inhibitors, such as cetuximab, in two *CRTC1-MAML2* fusion positive cells (FDR < 0.1%), mediated as a result of paracrine induction of EGFR signaling^20^ (Fig. 3c). Following manual curation, most associations between fusions and drug sensitivity could be readily explained by known interactions (n = 66; 30%), mutations in secondary genes (n = 7; 3%), and fusions that were either not in-frame (n = 77; 34%) or not seen in patient samples (n = 131; 57%). The remaining associations (n = 35; 15%) generally involve poorly described fusions present in 2 or 3 cell lines, making drug sensitivities difficult to interpret.

This analysis was limited in power by the small number of recurrent fusions genes in the dataset. Nonetheless, it suggests that besides well-established oncogenic fusions, there are few recurrent gene fusions that could be used as therapeutic biomarkers for repurposing of existing anti-cancer drugs. We did, however, observe potent drug sensitivity to particular drugs in individual cell lines with rare fusions.

### Functional analysis using CRISPR-Cas9 loss-of-fitness data

Our analysis of fusions using drug sensitivity data was limited by their low frequency and the limited number of targets covered by available drugs. Here, we complemented our fusion identification pipeline with CRISPR-Cas9 loss-of-fitness screens to systematically assess their functional implications. CRISPR screens typically target a gene with 5-10 single guide RNAs (sgRNA) and aggregate fold-changes to calculate gene-level depletion values. By contrast, we took advantage of individual sgRNA fold-changes to query the functional importance of gene regions. We assembled CRISPR whole-genome drop-out screening data from Project Score at the Sanger Institute (manuscript under review and can be made available upon request), Achilles project and Wang *et al.*, that together span 371 cell lines from 33 different cancer types^6,21^. We then mapped the coordinates of the sgRNAs targeting either of the fusion genes, and classified them as mapping or non-mapping sgRNAs, depending on whether they targeted the fusion transcript or not (Fig 4a). We calculated a fusion essentiality score (FES) for each gene fusion transcript partner as the differential scaled fold-change between mapping and non-mapping guides, or, where a fusion transcript had no non-mapping guides, the value of non-mapping guides was set to zero.

**Figure 4:**
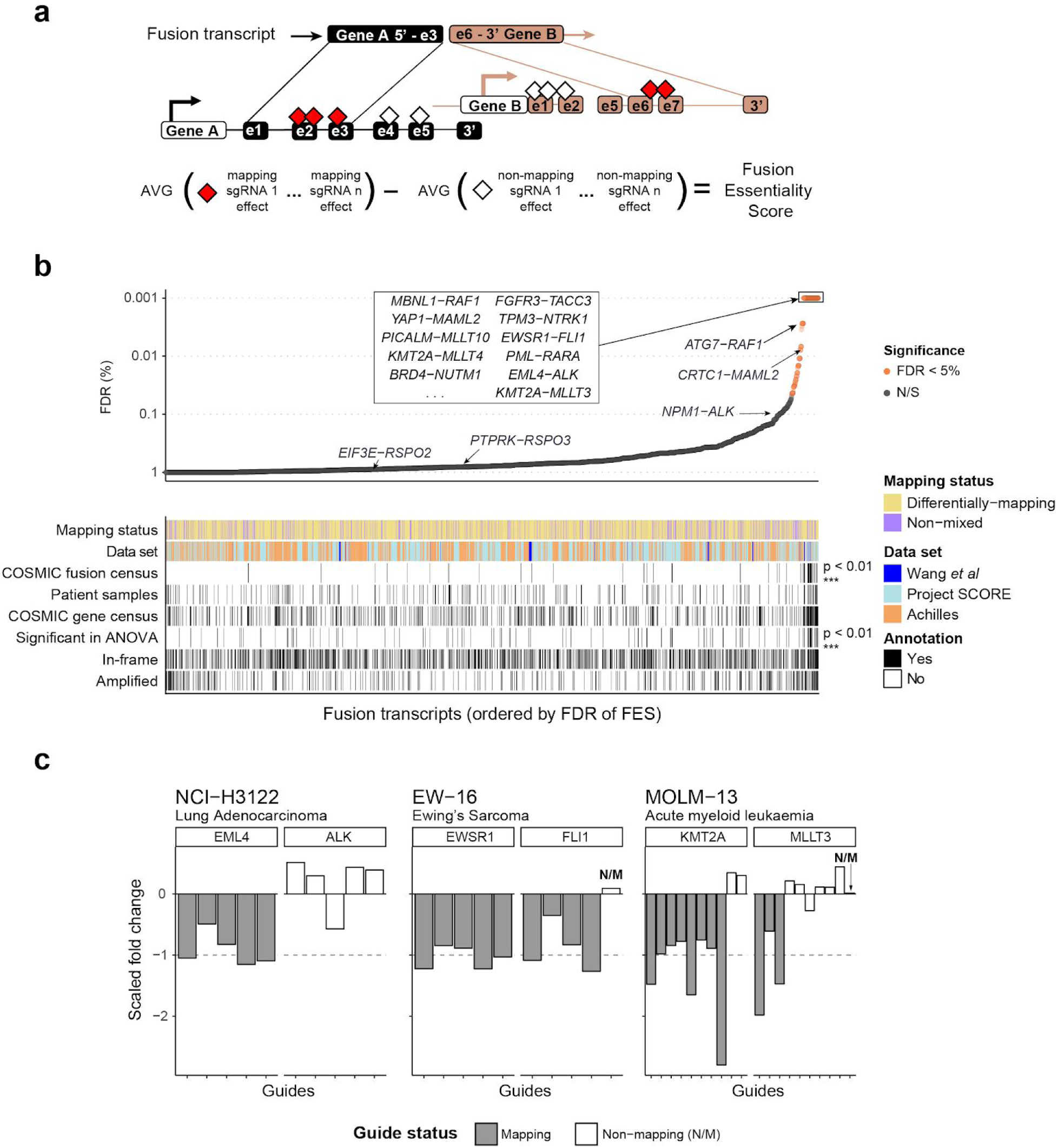
CRISPR whole-genome screening data identifies functional fusions. (a) Schematic of the analytical approach mapping individual sgRNAs to each gene in a fusion transcript and calculation of fusion essentiality scores (FES). (b) False discovery rate (FDR) of FES scores (see online methods) for all testable fusion transcripts (n = 2,821). Transcripts with at least one mapping and one non-mapping guide are “differentially mapping”, while transcripts with only mapping guides are “non-mixed”. Shown are whether fusions transcripts are listed in the COSMIC fusion census, found in patient samples^13^, if one of the partner genes is described in the COSMIC Gene Census, if one of the partner genes is amplified, and if the transcript is in-frame. Statistical significance of FES and false discovery rate (FDR) was based on 10,000 randomizations of data (see Online Methods). Selected known oncogenic fusions and other fusions of interest are highlighted. (c) Examples of functional fusion transcripts identified in specific cancer cell lines based on FES scoring. Each bar is the scaled fold-change of an individual sgRNA to fusion 5’ and 3’ end partner genes, and colored by fusion mapping or non-mapping sgRNA. Dotted line is at −1 (to which known essential guides were scaled, see methods). AVG = average. N/S = not significant at 5% FDR

We identified mapping sgRNA for 2,821 fusions transcripts, of which 129 fusion transcripts (5%) (representing 103 fusion events) were significantly associated with decreased cell fitness when targeted in at least one data set (Fig. 4b, Supplementary Table 8). Using a gene-set enrichment analysis, we found an enrichment in significant FES for fusions transcripts in the COSMIC fusion database of driver oncogenic fusions (p < 0.001), which included well-known oncogenic fusions like *EML4-ALK* (FDR < 0.5%), *EWSR1-FLI1* (FDR < 0.5%) and *KMT2A-MLLT3* (FDR < 0.5%) and TPM3-NTRK1 (FDR < 0.5%) (Fig 4b and c, Supplementary Fig. 3). Among the most significant associations were *YAP1-MAML2* fusions (FDR < 0.01%), *DDX6-FOXR1* (FDR < 0.5%) and *PICALM-MLLT10* (FDR < 0.5%). Interestingly, there was no enrichment in significant FES for fusion transcripts that were: (i) previously reported in patient samples; (ii) fusion transcripts that are in-frame vs. not in-frame; (iii) fusion transcripts in amplified regions, which are associated with non-specific fitness effects in CRISPR screens; (iv) nor for fusion transcripts that involve genes in the COSMIC Census^22^. These results demonstrate the ability to identify functional fusion transcripts using CRISPR-Cas9 screening datasets, but that for most tested fusions, including those identified in patients and involving cancer driver genes, we did not detect evidence supporting a functional role for cancer cell fitness.

### Function of oncogenic gene fusions across different histologies

Our analyses provide multiple lines of evidence supporting the functional role of gene fusions. Here we provide insights into the pathogenic role of specific gene fusions. Specifically, known oncogenic fusion are identified in alternative tissue types, pointing to strategies for repurposing clinically approved drugs in rare subsets of fusion-positive cancers. Furthermore, previously uncharacterised fusions are shown to drive aberrant signalling in cancer cells.

### Druggable RAF1 fusions in pancreatic adenocarcinoma cells

Rare *RAF1* fusions have been reported in patient tumors^23–25^, and *RAF* fusions are biomarkers of response to MAPK pathway inhibition. We identified an in-frame *ATG7-RAF1* fusion in PL18, a pancreatic adenocarcinoma cell line (Fig. 5a). The fusion was confirmed by Sanger sequencing across the breakpoint and fluorescence in situ hybridization (FISH) (Fig. 5b and Supplementary Fig. 4). The fusion removes the N-terminal regulatory regions but retains an intact *RAF1* protein kinase domain, suggesting it results in constitutive kinase activation.

**Figure 5:**
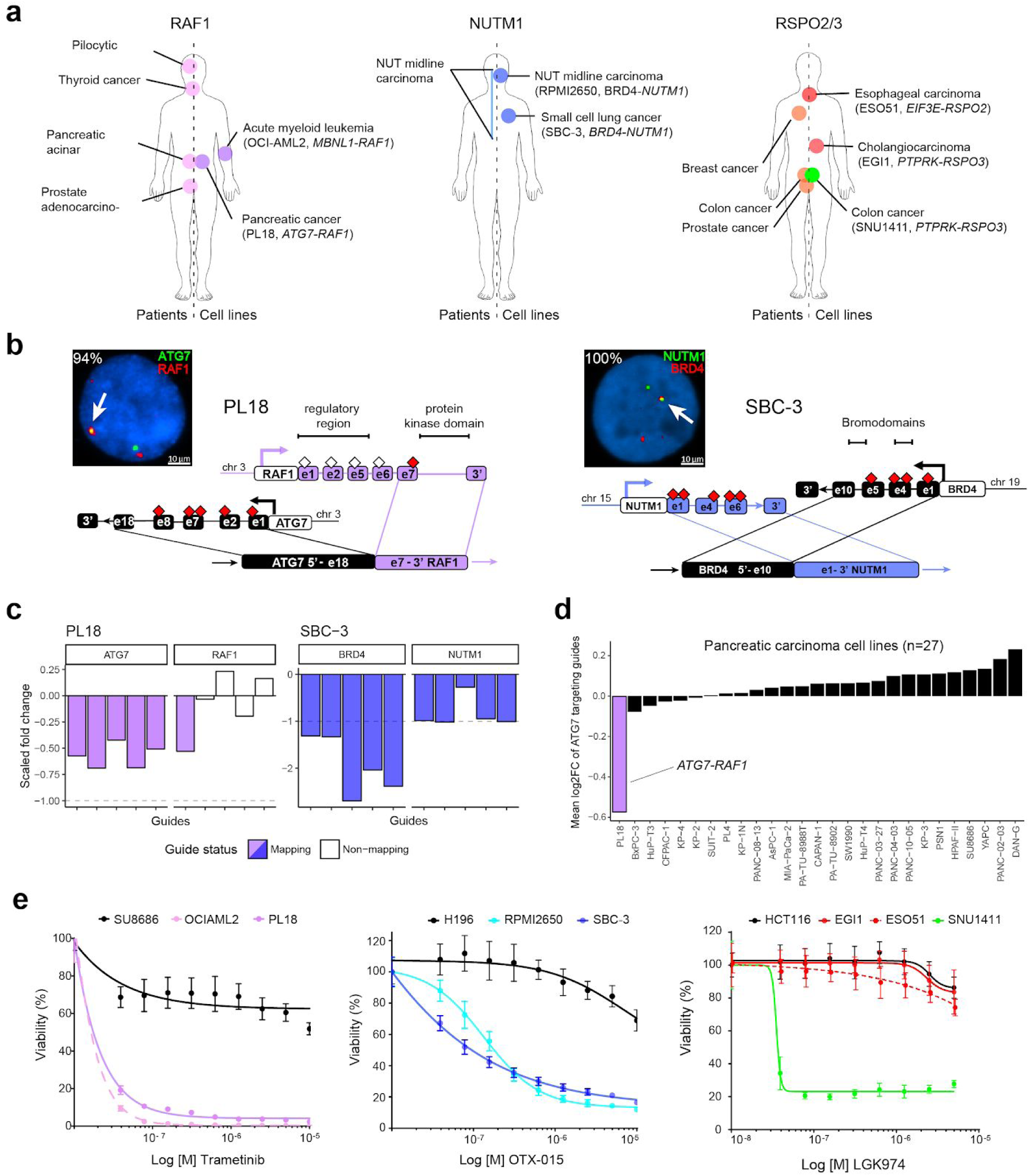
Therapeutically actionable oncogenic fusions identified across different histologies. (a) Evidence for oncogenic RAF1, NUTM1 and RSPO2/3 gene fusions identified in patients previously (left) and cell lines in this study (right). Cell lines carrying known oncogenic fusions used as positive controls for validation experiments are reported. (b) Interphase FISH of ATG7-RAF1 (left) and BRD-4-NUTM1 (right) gene fusions (arrows) in PL18 and SBC-3 cell lines. The percentage of fusion-positive interphases are reported in white text. Schematic representations of each fusions is also represented. Only exons involved in the breakpoint or displaying fusion mapping sgRNAs (red rectangles) or non-mapping sgRNAs (empty diamonds) are shown. (c) Fold change FES values of sgRNAs targeting ATG7 and RAF1 genes in PL18 (left) and BRD4 and NUTM1 genes in SBC-3 cells (right). Colored bars indicate values of sgRNAs targeting the exons involved in the fusions. (d) PL18 cells showed the highest depletion of fusion-targeting ATG7 guides across the entire panel of screened pancreatic cancer cell lines. (e) Viability assay on PL18, SBC-3, EGI1 and ESO51 cells treated with MEK (Trametinib), BET (OTX-015) and PORCN (LGK974) inhibitors, respectively. SU8686, H196 and HCT116 cells are pancreatic, small cell lung cancer and colorectal cancer negative controls. OCIAML2, RPMI2650 and SNU1411 are, respectively, a RAF1-rearranged leukemia, a NUTM1-rearranged NMC and a RSPO3-rearranged CRC cell line included as positive controls. Data are average ± SD of three technical replicates from one representative experiment.

Only mapping sgRNAs targeting the portion of the two genes involved in the fusion were significantly depleted, resulting in a significant FES (Fig. 5c). Moreover, of the 27 pancreatic cancer cell lines analysed by CRISPR screening, *ATG7* fusion-targeting sgRNA were only depleted in PL18 cells (Fig. 5d). Unlike >90% of pancreatic tumors and cell lines that have activating mutations in KRAS^8,26^, PL18 has a wild-type *KRAS* allele, but retained potent sensitivity to downstream MEK pathway inhibitors trametinib and PD0325901 (Fig. 5e and Supplementary Figure 4). An *ATG7-RAF1* rearrangement was previously reported in another KRAS-wt pancreatic cancer model^27^. Furthermore, we mined sequencing data for 126 pancreatic adenocarcinoma patient-derived xenograft (PDX) models and identified an additional *KRAS* wild-type tumor with a *PDZRN3-RAF1* fusion which conserves the *RAF1* kinase domain (Supplementary Figure 4). Our data support emerging evidence for rare recurrent and potentially therapeutically actionable *RAF1* rearrangements in *KRAS* wild-type pancreatic cancer.

### Druggable BRD4-NUTM1 fusion in lung cancer cells

*BRD4-NUTM1* fusions genetically define NUT midline carcinoma (NMC), a rare and aggressive neoplasm that usually arises in the midline of the body with marked sensitivity to BET bromodomain inhibitors (BETi)^28,29^. We identified a novel in-frame *BRD4-NUTM1* fusion in SBC-3, a small cell lung carcinoma (SCLC) cell line and confirmed the fusion by Sanger sequencing and FISH (Fig. 5a and b, Supplementary Fig. 4). Based on CRISPR data on 206 cell lines screened at Sanger, *NUTM1*-targeting guides were highly depleted only in SBC-3 cells, and the fusion was associated with a significant FES (Fig. 5c and Supplementary Figure 4). Moreover, SBC-3 cells displayed marked sensitivity to four different BETi (Fig. 5e and Supplementary Fig. 4). Unlike >95% of SCLC tumors and cell lines, SBC-3 cells do not have alterations in *RB1* or *TP53*, nor do they express SCLC-specific neuroendocrine markers such as CgA, NSE and synaptophysin (Supplementary Figure 4). The *BRD4-NUTM1* fusion was specifically associated with high NUTM1 transcript expression in cell lines (Fig. 2 and Supplementary Figure 4). Therefore, we mined TCGA expression data for SCLC and non-SCLC (NSCLC) searching for samples displaying high *NUTM1* mRNA levels. We identified a single NSCLC sample, displaying *NUTM1* mRNA outlier expression (Supplementary Figure 4) and carrying a *NSD3-NUTM1* rearrangement^13^ (Supplementary Fig. 4), a chimeric oncoprotein recently identified in NMC patients and previously associated to BETi sensitivity^30^. *NUT* rearrangements were identified in rare subpopulation of SCLC and NSCLC patients^31,32^ and recent studies established that *NUT*-associated fusions can occurs in tumors outside the midline, such as soft tissue, brain, and kidney^33^. Our preclinical data suggest that *NUTM1* fusions could represent an actionable driver event in lung cancer with immediate potential clinical implications.

### Lack of pathway dependence in cells with canonical R-spondin fusions

Aberrant expression of RSPO2/3 fusion transcripts synergize with WNT-ligands to trigger WNT pathway activation in APC wild type colorectal cancer (CRC)^14^. WNT pathway blockade with porcupine inhibitors is effective in *RSPO3*-rearranged CRC preclinical models^34,35^ and clinical trials in patients with *RSPO2/3*-fusion positive tumors of any histological origin are ongoing (NCT01351103). Here, we detected and validated two unreported canonical R-spondin fusions in cell lines derived from biliary tract (EGI-1; *PTPRK-RSPO3* fusion) and esophageal (ESO51; *EIF3E-RSPO2* fusion) cancer by PCR and FISH (Fig. 5a and Supplementary Fig. 5). Aberrant expression of *RSPO2/3* was detected in both cell lines (Fig. 2b and Supplementary Fig. 5). Similarly, through mining TCGA esophageal cancer data, we found that a tumor with high *RSPO3* expression (>95th percentile) was positive for a canonical *RSPO3* fusion^13^ (Supplementary Fig. 5). Surprisingly, sgRNA mapping to the fusion were not associated with significant FES, and EGI-1 and ESO51 were insensitive to WNT pathway blockade (Fig. 5e and Supplementary Fig. 5). This was in contrast to SNU1411, a positive control CRC cell line model addicted to WNT-pathway activation by rearranged *RSPO3*, which was sensitive to multiple porcupine inhibitors^36^ (Fig. 5e and Supplementary Fig. 5). No additional alterations in the WNT pathway were detected in these cell lines as a possible explanation for lack of sensitivity. Thus, our results with RSPO2/3 fusions point to an element of tissue specificity in mediating the functional role of some fusions and differences in drug response, with potentially important implications for repurposing of WNT pathway inhibitors across different RSPO-fusion positive tumour types.

### Recurrent YAP1-MAML2 fusions drive Hippo-pathway signalling in different tissue types

We next used our CRISPR screening data to investigate the function of poorly understood fusions. Recurrent *YAP1-MAML2* fusions were identified in AM-38 (glioblastoma), ES-2 (ovarian carcinoma) and SAS (head and neck carcinoma) cell lines (Fig. 6a). We validated the fusion events in all three cell lines by PCR, and interphase and fiber FISH (Fig. 6b and Supplementary Fig. 6). *YAP1-MAML2* fusions have been reported in nasopharyngeal carcinomas and in a sample from a patient with skin cancer^13,37^, but not in the 3 tumour types reported here (Fig. 6a). The fusion brings together exons 1-5 of *YAP1* and exons 2-5 of *MAML2*, a transcript structure that is conserved across all three cell lines and patient samples (Fig. 6c).

**Figure 6:**
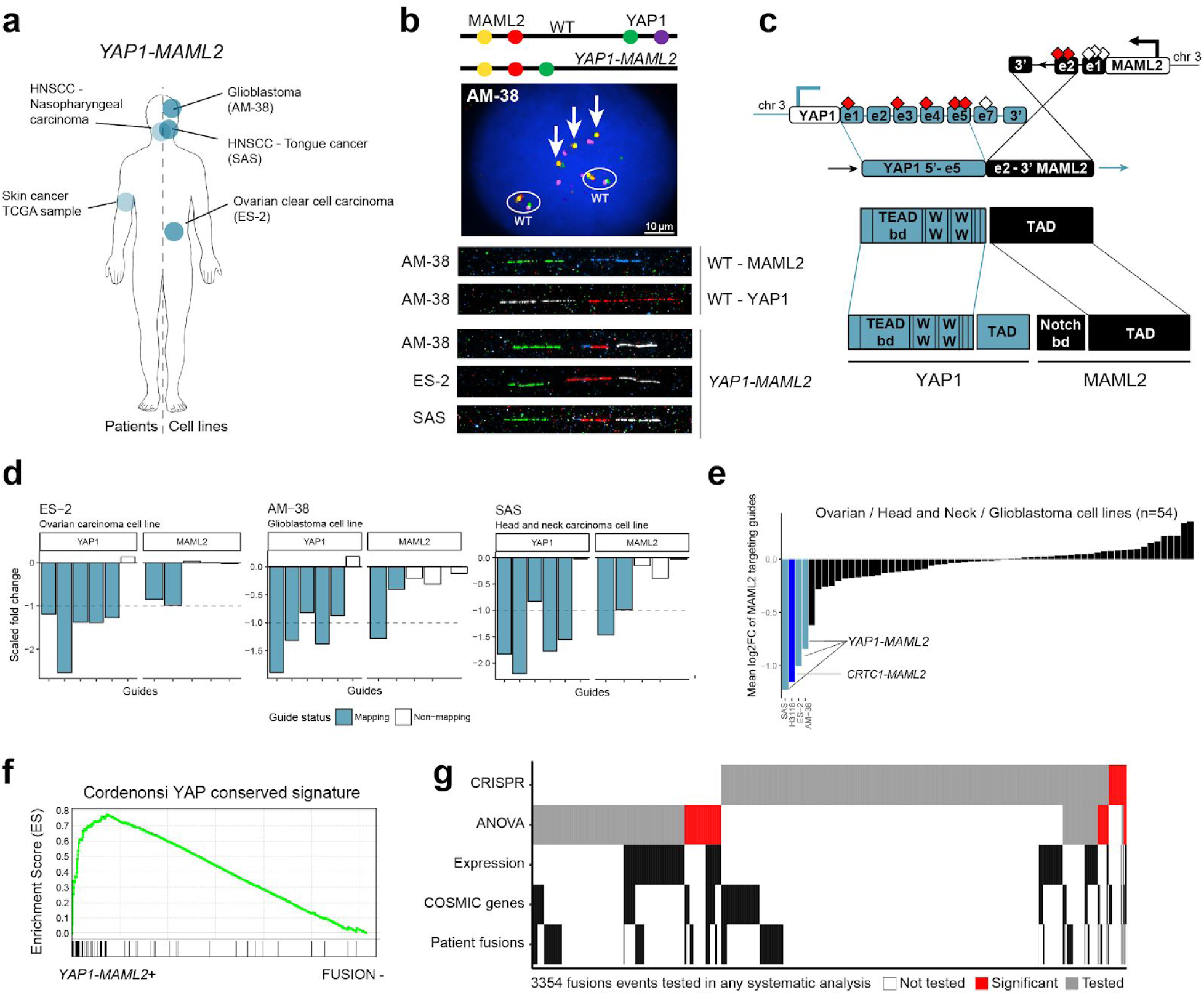
Recurrent YAP1-MAML2 gene fusions activate Hippo-pathway signalling. (a) YAP1-MAML2 gene fusions identified in patient tumor samples (left) and cell lines (right). (b) Interphase FISH (AM-38 only) and fiber FISH targeting YAP1-MAML2 fusion (arrows; cells are polyploid with wild-type chromosomes circled) in AM-38, ES-2 and SAS cell lines. Probes used and chromosomal position are shown schematically. YAP1 and MAML2 are both on chromosome 3. (c) Schematic representation of YAP1-MAML2. Only exons involved in the breakpoint or displaying fusion mapping sgRNAs (red diamonds) or non-mapping sgRNAs (empty diamonds) are shown. YAP1 and MAML2 functional domains involved in the fusion are indicated. (d) Fold-change values of sgRNAs targeting YAP1 and MAML2 genes in ES-2, AM-38 and SAS cell lines. Colored bars indicate the values of fusion mapping sgRNAs. (e) YAP1-MAML2 fusion-positive cell lines show the highest depletion of fusion-targeting MAML2 guides across ovary, head and neck, and glioblastoma cell lines (n= 54). Cell line in dark blue (H3118) harbors a known CRTC1-MAML2 fusion (Fig. 3c)^20^. (f) Gene set enrichment analysis (GSEA) of YAP1 gene signature in YAP1-MAML2 positive cells (n= 3) vs fusion negative cell lines (n= 1008). (g) Heatmap representing 3,345 fusion events tested in CRISPR and ANOVA systematic analyses. Significant associations are highlighted in red. Fusions events are annotated if one of the partner gene is significantly overexpressed using our linear regression model, contain a COSMIC cancer driver gene, or have been detected in patient samples.

A functional role for *YAP1-MAML2* fusions has not been previously been reported. We found that *YAP1-MAML2* fusions were significantly associated with decreased cell fitness when targeted in the CRISPR screen (Fig. 4b and 6d). Furthermore, loss of fitness in response to *MAML2*-depletion is unique to *MAML2*-fused cell lines in the three cancer types where the fusion is observed (Fig. 6e). YAP1 overexpression is linked with poor prognosis, chemoresistance and resistance to cell death in multiple solid tumors. YAP1 is a transcriptional co-activator of the Hippo pathway through binding with the TEAD1 transcription factor and MAML2 is a transcriptional co-activator involved in NOTCH signaling^38^. *YAP1-MAML2* fuses the transcriptional activation domain of MAML2 with the TEAD-binding domain of YAP1. Intriguingly, ES-2 and AM-38, although not SAS, also showed essentiality for *TEAD1* in the CRISPR-dropout screen (data not shown), suggesting that the fusion protein signals through TEAD1.

To further investigate fusion protein activity, we performed gene-set enrichment comparing the three *YAP1-MAML2* fusion positive cell lines against all others. Of 189 pathways tested, the YAP1-conserved signature was the most significant hit (adjusted p < 0.001) (Fig. 6f). The same signature was highly enriched when ES-2 was compared against all other ovarian cancer cell lines and SAS against all other head and neck cell lines, while expression of prototypic tissue-specific oncogenic signatures, such as estrogen receptor signaling in ovary, were depleted (Supplementary Fig. 6). In summary, we demonstrate that recurrent *YAP1-MAML2* fusion are associated with increased YAP1 signaling and required for cell fitness. Our results support targeting the Hippo-signalling cascade in *YAP1-MAML2* fusion-positive tumours.

## Discussion

Thousands of gene fusion transcripts have been reported from the analysis of large cohorts of tumor samples and preclinical models^13,39,40^. Most fusions are likely to be passenger events due to chromosomal instability or artifactual. Critically, although studies have investigated the functional role of a small number of individual fusions, to the best of our knowledge, there have been no comprehensive analyses to investigate the functional landscape of gene fusion across diverse tissue histology. We developed a multi-algorithm fusion calling pipeline, and integrated large-scale genomic and functional datasets, including CRISPR-Cas9 whole-genome screening data, to systematically identify gene fusions required for the fitness of cancer cells. Since fusions can have diagnostic, prognostic and therapeutic utility, our analysis could have clinical implications.

We demonstrate that most fusions are rare events occurring in a small number of cell lines, and their frequency and distribution broadly matches what is observed in patient tumours. In total, we tested 3,354 fusion events and found supporting evidence of a functional role for 368 (11.8%) by either CRISPR data (n = 103) or ANOVA analysis (n = 284)(Fig. 6g). Of those, 142 (38.5%) involved a COSMIC gene, 58 (16%) were listed in the COSMIC fusion census, and 107 (29%) were also called in TCGA patient samples. Thus, many fusions with supporting functional evidence are poorly understood and do not contain known driver genes, suggesting that there are gaps in our knowledge of genes with roles in cancer cell fitness. Conversely, most fusions tested were not required for cancer cell fitness, including fusions with known cancer driver genes. Collectively, we find that most fusions tested do not have supporting functional evidence, emphasizing the importance of analyses to ascribe function when interpreting fusions identified using genomic sequencing.

We cannot exclude the possibility that some fusions are required for aspects of the malignant phenotype or carcinogenesis not measured here, such as tumor initiation, paracrine signaling, host–tumor cell interaction and metastasis. Our CRISPR-based approach was not suitable for testing fusions which did not have mapping sgRNA. Moreover, it only captures fusions which induce gain-of-function or dominant-negative effects, but is not able to identify recessive effects such as inactivation of a tumour suppressor.

Gene fusions are used as therapeutic biomarkers to enrol patients in clinical trials and to direct clinical care, often in diverse histologies and clinico-pathologic subtypes. Notable examples are NTRK and ALK fusions, originally identified as effective biomarkers of response to targeted agents in NSCLC patients and occurring at low frequencies (<1%) in a variety of malignancies^41–43^. We provide specific and previously undescribed data on fusions involving *RAF1, ROS1* and *BRD4* that suggest existing drugs could be repurposed for use in rare pancreatic, breast, and lung cancers. Further studies using tumour xenograft models would support the *in vivo* efficacy of these findings and could pave the way for their clinical application. More broadly, these results support the use of validated oncogenic fusions as therapeutic biomarkers in diverse histologies, and the utility of basket trials for clinical development of drugs targeting fusion proteins irrespective of tumour type, such as the type used for the development Entrectinib in solid tumour with *ALK, ROS1* and *NTRK* fusions^43^.

A notable exception in our analysis was the differential sensitivity to WNT-pathway inhibition of CRC versus biliary tract and esophageal cancer cell lines with canonical R-spondin fusions. This suggests that tissue context could impact the functional role of some fusions as has been observed for oncogenes (e.g. *BRAF*-mutated CRC^44^), with implications for development of genotype-directed trials in multiple tissue histology. Further investigations are warranted to understand this difference, and drug combinations could be evaluated in these specific context to overcome resistance similar to what is in clinical development for *BRAF*-mutated CRC^44^.

We identified and functionally evaluated less well studied gene fusions, as exemplified by *YAP1-MAML2* rearrangements, which are required for cell fitness in multiple histology and associated with increased YAP1 signaling. Given the emerging role of YAP1/TEAD1 and the Hippo pathway in cancer, there is interest in pharmacological inhibition of Hippo-signaling as an anticancer therapeutic strategy^45^. We provide preclinical evidence supporting inhibition of this signaling axis in *YAP1-MAML2* fusion positive tumors, with could pave the way for clinical development in a rare but defined patient population.

Our analysis supports the use of functional perturbation studies in preclinical models as an unbiased platform to systematically assess the impact of fusions in cancer. Extending this approach to a larger set of cancer models that represents the histopathologic and genomic diversity of patient tumours could reveal additional new insights with clinical relevance. In conclusion, we find that most fusions are not functional, with important implication for the interpretation of tumour sequencing data. Nonetheless, we identified fusion gene drivers of carcinogenesis which could represent future targets for drug development and specific actionable leads with potential for immediate clinical development in defined fusion-positive patients.

## Author contributions

The project was conceptualised by MG, GP, EC, GB and UM. Majority of molecular biological experiments were designed by GP and EC and performed by GP. Majority of computational analyses designed by EC and GP and implemented, data curated and processed by EC. RNA-Seq data pre-processing was performed by GB, AM and AB. Gene expression analysis was performed by LA and JS-R. Gene set enrichment analysis was performed by GP. CRISPR-Cas9 screening data access and related advice was provided by FB, FI, EG, ES and KY. Statistical and computational advice provided by EG and FI. PCR validations performed by EA. FISH experiments were performed by BF, RB and FY. The manuscript was prepared and written by MG, GP and EC. All authors edited and approved the manuscript. Funding was acquired by MG and JS-R. Project administration by MG.

### Acknowledgments

We thank Sandra Louzada for support with DNA fibres probe preparation.

## Data and materials availability

All data is available in the main text or the supplementary materials

## Supplementary Materials

Methods, Supplementary Fig. 1-6, Supplementary Tables 1-10

