## Supplementary Material for "Functional linkage of gene fusions to cancer cell fitness assessed by pharmacological and CRISPR/Cas9 screening"

### Supplementary Tables

**Supplementary Table 1:** Annotation of 1,034 human cancer cell lines used in our study and the source of RNA-seq data. All cell lines are part of the GDSC cancer cell line project (see COSMIC IDs).

**Supplementary Table 2:** List and annotation of 10,514 fusion transcripts found in 1,011 cell lines.

**Supplementary Table 3:** Significant results from differential gene expression for recurrent fusions.

**Supplementary Table 4:** Annotation of gene fusions for aberrant expression of the 3 prime end gene.

**Supplementary Table 5:** High-throughput cell line drug sensitivity data (IC50's) used in this study together with compound annotation. Column names are COSMIC id's of cell lines.

**Supplementary Table 6:** Significant results from the drug-association analysis using cancer functional events.

**Supplementary Table 7:** Significant results from the drug-association analysis using gene fusions.

**Supplementary Table 8:** Fusion essentiality score and significance calculation for all 2,821 fusion transcripts with mapping guides. For 525 fusion transcripts where multiple data sets contained mapping guides, both are reported.

**Supplementary Table 9:** List of primers sequences used to validate gene fusions.

**Supplementary Table 10:** List of BAC and fosmid clones used in the FISH validation.

### Methods

#### Sample selection

A collection of 1,011 cancer cell lines have been compiled from publicly available repositories as well as private collections and maintained following supplier guidelines. These cell lines were selected to be genetically unique based on STR and SNP fingerprints (http://cancer.sanger.ac.uk/cell_lines/download). STR profiles matched those in public repositories or match published STR profiles. The cells lines have been extensively characterized using whole exome sequencing (EGAD00001001039), Affymetrix SNP6 based copy number analysis and genotyping (EGAD00010000644). All data are available from the COSMIC Cell Line Project (COSMIC-CLP) (https://cancer.sanger.ac.uk/cell_lines). Comparing SNP6 genotyping data and somatic variant data we verified that COSMIC-CLP overlap extensively with those characterised by the Cancer Cell Line Encyclopaedia (TCGA). See Supplementary Table 1 for a complete description of the cell lines and their molecular annotation.

#### RNAseq data acquisition and identification of fusion transcripts

RNA-seq data for 589 cell lines was obtained from the Cancer Genome Hub (CGHub) and 450 cell lines were sequenced at the Sanger Institute (EGAS00001000828). For 23 cell lines, sequence was obtained from both CGHub and Sanger Institute to allow comparison of the output based on sequence from the two studies. Where replicated data sets were available, we took forward only fusions called from Sanger Institute sequencing data for our final analysis.

For sequencing performed at the Sanger Institute, cell line pellets were collected during exponential growth in RPMI or DMEM/F12 and were lysed with TRIzol (Life Technologies) and stored at -70°C. Following chloroform extraction total RNA was isolated using the RNeasy Mini Kit (Qiagen). DNAse digestion was followed by the RNAClean Kit (Agencourt Bioscience). RNA integrity was confirmed on a Bioanalyzer 2100 (Agilent Technologies) prior to labelling using 3' IVT Express (Affymetrix). Sequence libraries were prepared in an automated fashion on the Agilent Bravo platform using the stranded mRNA library prep kit from KAPA Biosystems. Processing steps were unchanged from those specified in the KAPA manual except for use of an in house indexing set. Three publically available gene fusion detection algorithms were used (TopHat-Fusion (v2.1.0), STAR-Fusion (v2.5.0) and deFuse (v0.7.0)) as described in GitHub (https://github.com/cancerit/cgpRna/blob/dev/README.md).

#### Fusion Transcript Filtering criteria

From the output of the three distinct fusion detection algorithms, we selected for analysis only fusions that were called with 4 or more reads that align directly across the breakpoint. We also required fusions to be called by at least two different algorithms. Next, we removed fusions identified from the analysis of 245 non-neoplastic samples downloaded from GTEx[1].

#### Fusion annotations

The frame of fusion transcripts was predicted using the GRASS algorithm that is built into the fusion-calling pipeline (https://github.com/cancerit/cgpRna/blob/dev/README.md / https://github.com/cancerit/grass). A list of known cancer driver genes was obtained from the COSMIC cancer census (https://cancer.sanger.ac.uk/cosmic/curation). Known cancer fusions, as well as annotation of tumour suppressor genes and oncogenes, were downloaded from the COSMIC fusion census (https://cancer.sanger.ac.uk/cosmic/fusion).

Our reference set of fusions identified previously in patient samples comes from an analysis of fusions in over 9,000 TCGA samples[2].

#### PCR validation of fusion transcript

cDNA was prepared using the Superscript double-stranded cDNA synthesis kit (Invitrogen) followed by SPRI clean-up. The cDNA was then subjected to ‘whole genome amplification’ using the Illustra GenomiPhi HY DNA Amplification Kit as per the manufacturer’s instructions. This WGA’ed cDNA was used as a template for the PCR validation. Generally two distinct sets of PCR primers were designed using Primer3 (http://www.bioinformatics.nl/cgi-bin/primer3plus/primer3plus.cgi) for each fusion junction tested (Primer sequences are in Supplementary Table 9). The primers were then checked by ePCR (http://www.ncbi.nlm.nih.gov/sutils/e-pcr/reverse.cgi?taxid=9606&db=2&orgdb=118&margin=200&mism=0&gaps=0) against the genome and transcriptome to make sure that they would not produce a PCR product of <5kb. PCRs were carried out in duplicate using two PCR programs: 1) 30 cycles, 95°C for 30s, 60 °C for 30s and 72 °C for 30 s; 2) a touchdown program reducing annealing temperature by 2 degrees every 2 cycles, dropping from 60 °C to 50 °C over ten cycles, with a final 20 cycles at 50 °C. For all cycles the melting and extension temperatures were 95 °C and 72 °C, respectively, and all stages were maintained for 30 seconds. Finally, the PCRs were checked by gel electrophoresis to confirm the presence of a product of the predicted size. To validate candidate fusions, PCR products were further checked by PCR product Sanger sequencing. In these cases the PCR products were first cleaned up using ExoSAP (Affymetrix) and then capillary sequenced by Eurofins Genomics (Ebersberg, Germany).

#### Differential fusion frequencies across cancer types

We designed an empirical permutation approach to identify genes with differential fusion frequencies across cancer types. Briefly, we built a binary matrix (genes x samples) where 1 indicates that the sample contains at least one fusion involving a gene G and 0 indicates that the gene G is not fused. Under the null hypothesis that gene alterations distribute homogeneously across cancer types (i.e., any sample from any cancer type has the same likelihood of having the gene G fused), we permuted 10,000 times the initially observed matrix using the algorithm BiRewire implemented in an R package[3]. This algorithm generates randomized binary networks that preserve marginal totals. Randomized networks are used to build a null distribution, modeling the likelihood of observing a gene fused in X samples. Thus, we can derive a *p*-value per gene and cancer type representing the probability of observing ≥ N samples with the gene G fused in the null distribution. Nominal *p*-values are adjusted using the FDR method.

#### Gene expression and differential gene expression analysis

Read counts per gene, based on the union of all exons from all possible transcripts, were used to calculate Reads Per Kilobase per Million (RPKM) as described previously[4]. To identify genes which expression is significantly altered when fused, we used a multiple linear regression approach. For each fusion-associated gene, the expression values *G* ($\log_2$ RPKM) from each sample *S* are modeled as a function of the fusion status of the gene in the sample *S* ($X_{fusion}$), a series of dependent covariates ($X_{covariates}$, including cancer-type in pan-cancer analyses, microsatellite instability status, and gene absolute CNA status), and a noise term (ψ):

$$G = \beta_{covariates} X_{covariates} + \beta_{fusion} X_{fusion} + \psi$$

The association between the fusion status and gene expression was defined by the regression coefficient ($\beta_{fusion}$) estimated with a multiple linear least squares regression. Significance of the regressors was estimated with a type-II ANOVA method from the car R package. For each cancer-type, *p*-values were adjusted for multiple testing correction using the Benjamini–Hochberg method. To annotate fusion genes for overexpression of the 3' gene, we select fusions where the expression level of the 3' gene is above the 95$^{th}$ percentile (i.e. 95% of the cell lines have an expression level lower than the one observed in the cell line carrying the fusion). Z-score RNA-seq values for TCGA samples from Esophageal Adenocarcinoma (187 samples) and Lung Squamous Cell Carcinoma (187 samples) were downloaded from cBioPortal[5].

#### Gene set enrichment analysis

Data for gene set enrichment analysis and expression of neuroendocrine markers for GSEA analysis, RNA-seq voom transformed gene expression measurements[6] were obtained from[4]. GSEA software was downloaded from the Broad Institute GSEA portal (http://software.broadinstitute.org/gsea/index.jsp) and applied, using default parameters and exploiting signal-to-noise metric for gene ranking. The significance of enrichment was estimated using 1,000 gene permutations. Heatmap of neuroendocrine genes in small cell lung cancer cell lines was generated by the GEDAS software 1.1.6 Beta[7].

#### Cancer functional event drug-association analysis

Potential confounding factors in our fusion–drug association analysis were identified by implementing a systematic analysis of associations between cancer functional events published by Iorio *et al*[8] for our panel of cell lines and our set of 409 drugs. The 717 cancer functional events used in the analysis included 281 genes with somatic coding mutations,

424 copy-number altered chromosomal segments and methylation status for 12 segments that included any gene altered by point mutations. The analysis was conducted as described in Garnett *et al.*[9] although our analysis only considered tissue type and MSI-status as covariates.

We identify 101 large-effect size significant cancer functional event–drug associations across 73 drugs when implementing the same cut-offs as in Iorio *et al.*[8] (FDR < 25%, p-value < 0.001, and positive and negative Glass Deltas > 1) reported in Supplementary Table 6.

#### Gene fusion drug-association analysis

The fusion ANOVA model was constructed as per Garnett *et al.*[9], but also includes as covariate any cancer functional event that was involved in a significant large-effect size association with a given drug. High-throughput drug sensitivity data were generated by the Genomics of Drug Sensitivity in Cancer (GDSC) project (www.cancerRxgene.org) at the Sanger Institute as previously described[8]. Details of compounds screened and cell lines sensitivity data are provided in Supplementary Table 5.

#### CRISPR screening data analysis

Single guide RNA (sgRNA) raw counts and guide annotations were obtained for 206 cell lines from Project SCORE (manuscript under review - note to reviewers: this manuscript can be made available if requested) performed at the Sanger Institute, 274 cell lines screened as part of Project Achilles by the Broad Institute[10], and 14 acute myeloid cell lines screened by Wang *et al*[11]. sgRNA target positions were converted from GrCh37 to GrCh38 using the NCBI remapper tool (https://www.ncbi.nlm.nih.gov/genome/tools/remap).

sgRNA raw counts were converted into log fold-changes and corrected for CRISPR-biases using CRISPRcleanR (https://www.biorxiv.org/content/early/2017/12/03/228189 ; https://github.com/francescojm/CRISPRcleanR). Corrected log fold-changes were scaled to essentials using the scale-to-essentials R function in the CERES package (https://github.com/cancerdatasci/ceres/blob/master/R/scale_to_essentials.R).

Positions of sgRNA were mapped onto fusion transcripts using a bespoke R script that takes into consideration: (1) the fusion transcript breakpoint and (2) the mapping location of the guides. Essentially, sgRNA map onto the 5' gene if the gene is on the positive DNA strand and the guides maps before the position of the fusion breakpoint, or if the gene is on the negative DNA strand and the sgRNA maps after fusion breakpoint. Guides map onto the 3' gene if the opposite is true.

In order to calculate a fusion essentiality score, we took the following steps: 1) Z-normalized the scaled and corrected log fold-changes across all cell lines screened for a given sgRNA; 2) subtracted the mean Z-score of non-mapping from that of the mapping sgRNA for both genes involved in a fusion transcript. Where a gene has no non-mapping sgRNA, the difference was taken from zero; 3) values obtained in step 2 for each gene were averaged to produce the fusion essentiality score (FES).

To assign statistical significance for the FES, we performed 10,000 randomizations of all scaled and corrected log fold-changes within cell lines. Randomized-FES were then calculated for each fusion transcript, as described in the above paragraph. A *p*-value of statistical significance was assigned to each fusion transcript as the fraction of randomized-FES that have a higher FES score than the non-randomized FES. For multiple-hypothesis correction, we calculated a false discovery rate for each *p*-value. Calculation of the FES and the subsequent randomization was performed independently for each data resource, since the sgRNA libraries were different in all cases.

#### Cell viability assay

Cell lines were seeded at different densities (1 – $3x10^3$ cells per well) in 100 µl complete growth medium in 96-well plastic culture plates at day 0. The following day, serial dilutions of drug were added to the cells in an additional 50 µl of medium. Plates were incubated at 37 °C in 5% $CO_2$ for five days, after which the cell viability was assessed by measuring ATP content through Cell Titer-Glo Luminescent Cell Viability assay (Promega). Luminescence was measured by Envision Multiplate Reader at day 7. Crystal violet growth assay were performed seeding 30 - 50 x $10^3$ cells in 6-well plates. After 24 hours, medium was replaced adding drugs as indicated. After 7-10 days of treatments cells were fixed with a solution of 3% paraformaldehyde and then stained with 0.05% crystal violet in distilled water.

#### PDX database

Gene fusion data for 126 pancreatic adenocarcinoma PDX models were downloaded from HuBase database (https://hubase2.crownbio.com; Crown Bioscience International, Santa Clara, California, United States).

#### Interphase and fiber FISH

Metaphase chromosomes were prepared from cell lines listed in Supplementary Table 10 using a standard method. Briefly, colcemid (Thermo Fisher Scientific) was added to a final concentration of 0.1 mg/ml for 1 h, followed by treatment with hypotonic buffer (0.4% KCl in 10 mM HEPES, pH7.4) for 10 min and subsequent fixation using 3:1 (v/v) methanol:acetic acid. Human fosmid and bacterial artificial chromosome (BAC) clones containing the genes of interest (Supplementary Table 10) list of BAC and fosmid clones used in the FISH validation) were provided by the clone archive team of the Wellcome

Sanger Institute. Probes were generated by whole-genome amplification with GenomePlex Whole Genome Amplification Kits (Sigma-Aldrich), from purified BAC fosmid DNA as described previously[12]. For interphase- and metaphase-FISH, probes were labeled directly with Atto488-XX-, Cy3-XX-, Texas Red-12- and Cy5-XX-dUTPs (Jena Bioscience), respectively. Slides pre-treatment included a 10 min fixation in acetone (Sigma-Aldrich), followed by baking at 65°C for 1 hour. Metaphase spreads on slides were denatured by immersion in an alkaline denaturation solution (0.5 M NaOH, 1.0 M NaCl) for 7-10 min, followed by rinsing in 1M Tris-HCl (pH 7.4) solution for 3 min, 1× PBS for 3 min and dehydration through a 70%, 90% and 100% ethanol series. The probe mix was denatured at 65°C for 10 min before being applied onto the denatured slides. Hybridization was performed at 37°C overnight. The post-hybridization washes included a 5 min stringent wash in 1× SSC at 73-75°C, followed by a 5 min rinse in 2× SSC containing 0.05% Tween®20 (VWR) and a 2 min rinse in 1× PBS, both at room temperature. Finally, slides were mounted with SlowFade Gold® mounting solution containing 4'6-diamidino-2-phenylindole (Thermo Fisher Scientific).

Indirectly labelled probes were used in Fiber-FISH with single-molecule DNA fibers. The preparation of single-molecule DNA fibers by molecular combing and fiber-FISH followed Louzada et al. (2017, Springer Protocols). The three-colour probe set was labelled with biotin-16-, DNP-11-, and digoxenin-11-dUTPs (Jena Bioscience), respectively, and visualized with Cy3-, FITC- and Texas-red conjugated antibodies, with the exception of post-hybridization washes, which consisted of three 5 min washes in 2× SSC at 42°C, instead of two 20 min washes in 50% formamide/50% 2× SSC at room temperature. Slides were examined using AxioImager D1 microscope equipped with appropriate narrow-band pass filters for DAPI, Aqua, FITC, Cy3, Texas red and Cy5 fluorescence. Digital images capture and processing were carried out using the SmartCapture software (Digital Scientific

UK). Ten randomly selected metaphase cells were karyotyped based on the M-FISH and DAPI-banding patterns using the SmartType Karyotyper software (Digital Scientific UK).

### Supplementary Figures

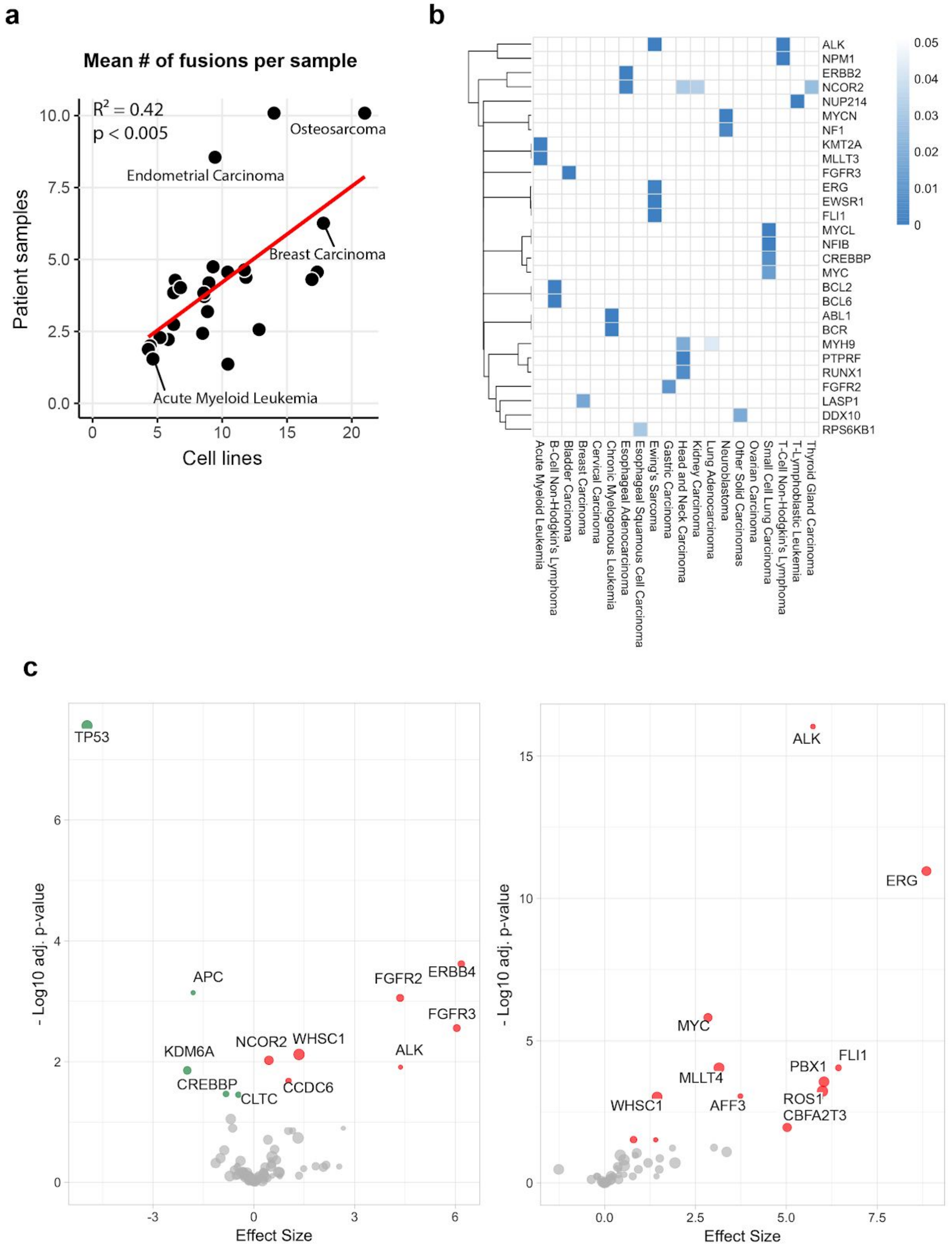

***Supplementary Figure 1: Gene fusions detected in cell lines and tumour and their impact on gene expression.*** *(a) Mean number of fusion events per sample across different cancer types was significantly correlated between cell lines and patient tumours[2]. Equivalent tissue annotations were present for 27 of 42 cancer types, covering 6,369 fusion events in 704 cell lines and 24,494 fusions events in 5,687 patient samples. (b) Heatmap for enrichment of fused genes within cancer types. Rows are genes and columns are cancer types. Colour intensity represents the strength of the association (adj. p-value). Only interactions with adj. p-value < 0.05 are represented. (c) Volcano plots of genes whose expression is significantly altered when at 5 prime end (left panel) and at the 3 prime end (right panel) of fusions. Only cancer driver genes are represented. X axis represents adjusted p-value using Benjamini & Hochberg (FDR) correction, while Y axis represents signed Cohen's d effect size. Circle size is proportional to the number of samples with the gene fused. Colours represent up-regulated (red) and down-regulated (green) genes, respectively. Labels indicate the name of the driver gene.*

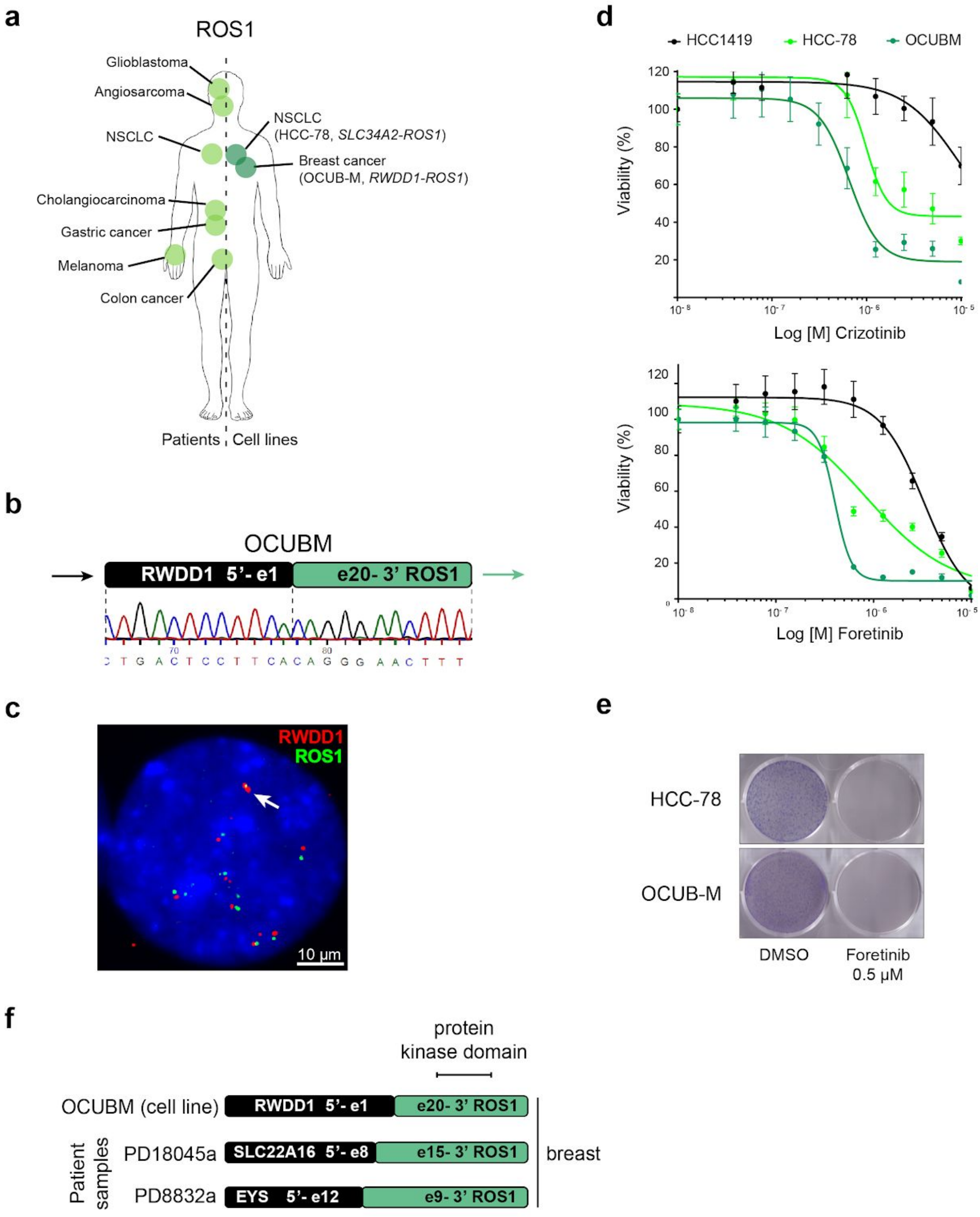


***Supplementary Figure 2: Validation of ROS1 fusion breast cancer cells.*** *(a) Evidence for oncogenic ROS1 gene fusions identified in patients previously (left) and cell lines in this study (right). HCC-78, a ROS1-rearranged non-small cell lung cancer (NSCLC) used as*

*positive controls for validation experiments is also reported. (b) Sanger sequencing across fusion breakpoint for RWDD1-ROS1 in OCUBM. (c) Interphase FISH images confirmed the presence of the RWDD1-ROS1 fusion in OCUBM cells. (d) Cell viability assays of OCUBM cells treated with ALK/ROS-inhibitors crizotinib (upper) and foretinib (lower). HCC-78 is a ROS1-rearranged non-small cell lung cancer (NSCLC) and HCC1419 is a fusion-negative control breast carcinoma cell line. (e) Colony formation assays of OCUBM and HCC-78 cells treated with foretinib. (e) Breakpoints of ROS1-fusions in cell lines and cancer patient samples.*

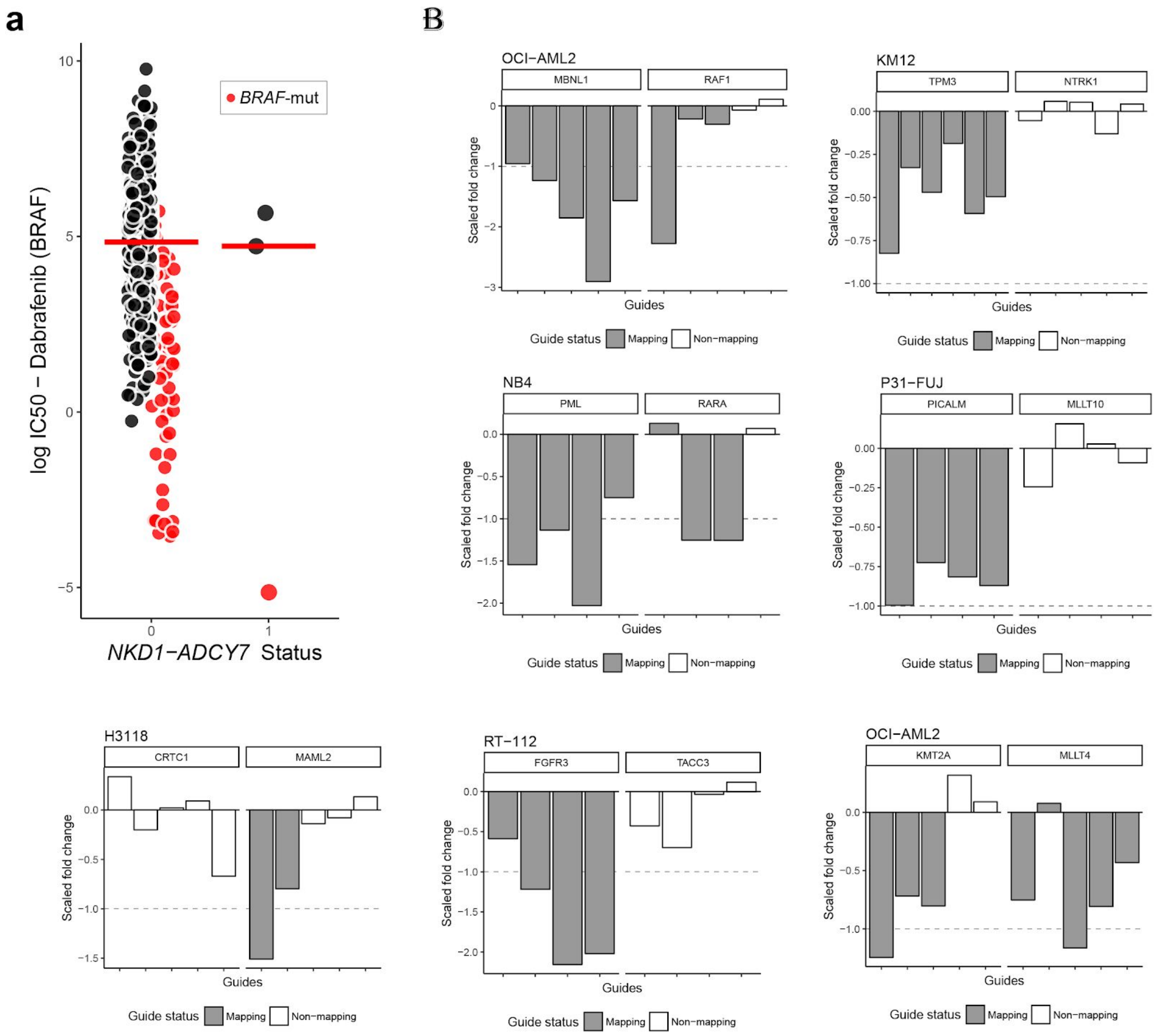


***Supplementary Figure 3: Example of a false-positive associations in ANOVA analysis and examples of significant fusion essentiality score for known oncogenic fusions.*** *(a) Using cancer events as covariate in our fusion-drug ANOVA successfully excluded false positive associations with fusion events. The association of a NKD1-ADCY7 fusion with BRAF inhibitor Dabrafenib is confounded by a BRAF-mutation in one of the cell lines. This association was no longer significant after implementing the covariates. (b) Examples of 7 known oncogenic fusions with a statistically significant fusion essentiality score using CRISPR-Cas9 loss-of-fitness data.*

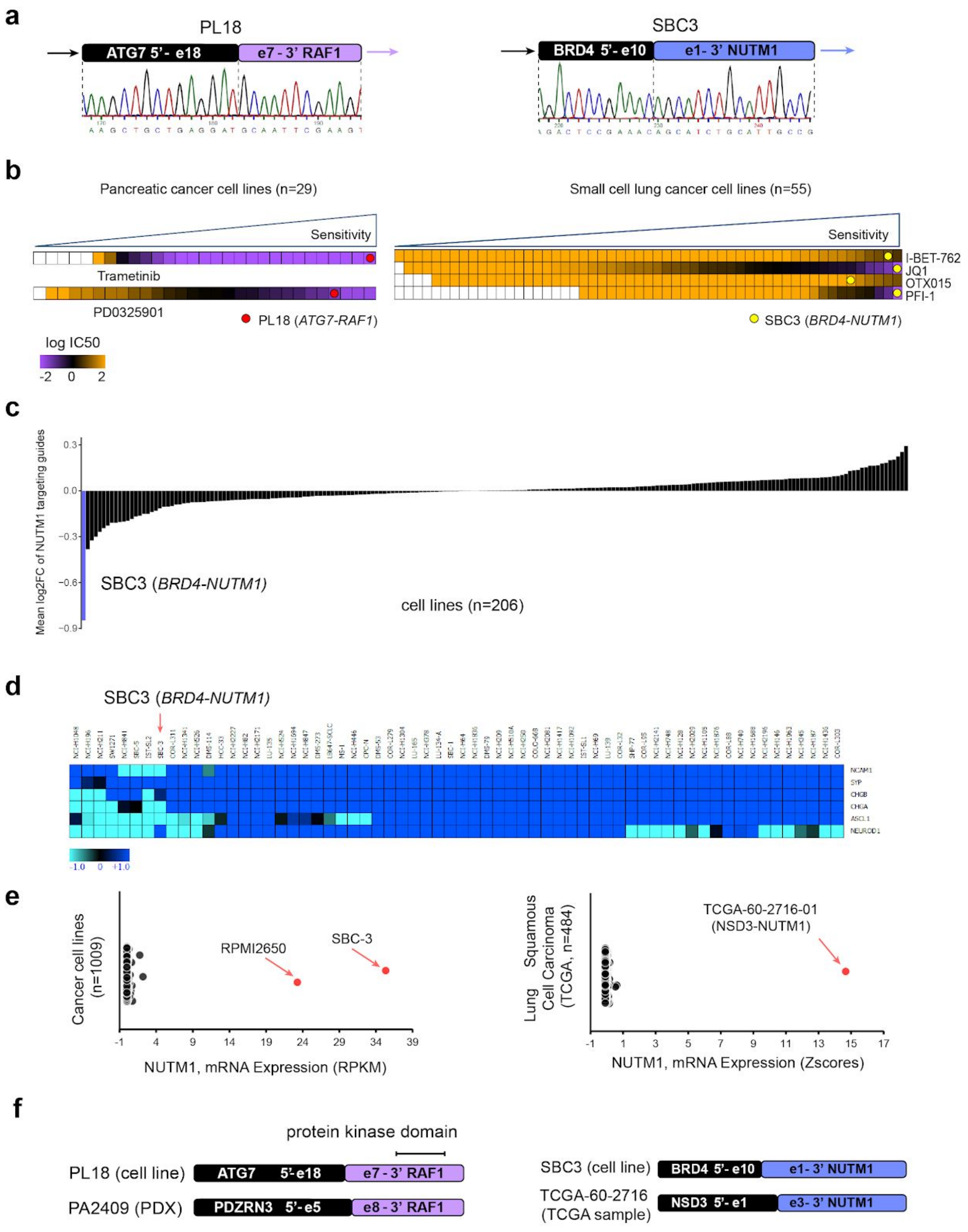


***Supplementary Figure 4: Validation of RAF1 and NUTM1 rearrangements in pancreatic and small cell lung cancer cells.*** *(a) Sanger sequencing across fusion breakpoints for ATG7-RAF1 in PL18 (pancreatic cell line) and BRD4-NUTM1 in SBC-3 (small cell lung*

*cancer cell line). (b) Heatmaps represents ranked trametinib and PD0325901 log IC50 values in pancreatic cancer cell lines (left) and BET inhibitors log IC50 values in small cell lung cancer cell lines (right) measured by GDSC high-throughput drug screening. PL18 cells are highly sensitive to both MEK inhibitors, SBC-3 cells are highly sensitive to multiple BET inhibitors. (c) Across 206 cell lines screened with the Sanger human CRISPR library v1, SBC3 has the highest depletion of NUTM1 fusion-targeting guides. (d) Expression of neuroendocrine markers across 64 small cell lung cancer cell lines. Unlike the majority of small cell lung cancer cell lines, SBC-3 shows low expression of typical neuroendocrine markers. (e) BRD4-NUTM1 fused cell lines (SBC3 and positive control cell line RPMI2650) show exceptionally high expression of NUTM1 compared to all other cancer cell lines (left panel). We identified a lung squamous cell carcinoma TCGA tumour sample with high NUTM1 expression and a NSD3-NUTM1 fusion (right panel). Z-score of RNA-seq values for TCGA samples were downloaded from cBioPortal. (f) Fusion breakpoints of RAF1 and NUTM1 rearrangements in cell lines/PDX models and patient samples. RPKM, Reads Per Kilobase Million.*

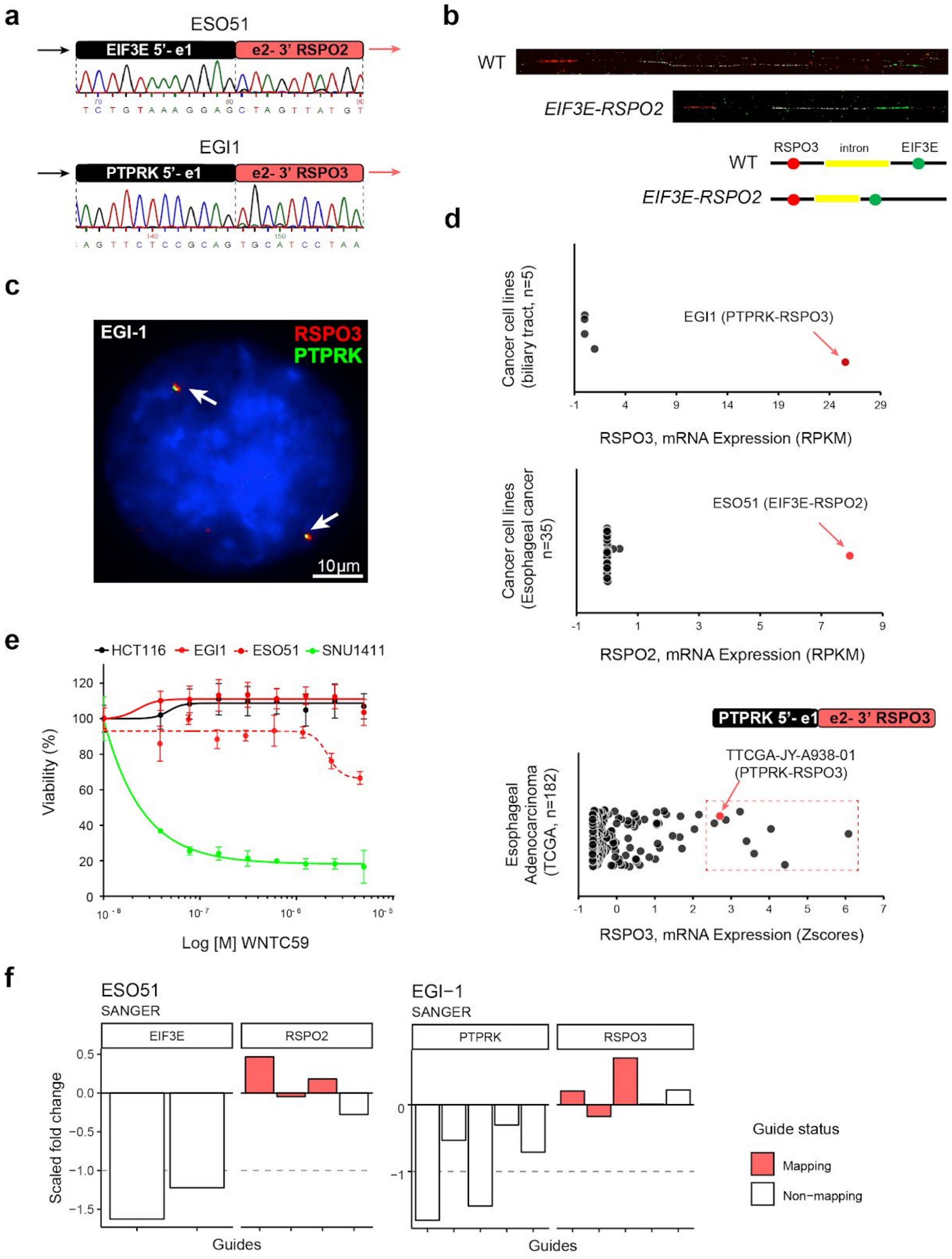


***Supplementary Figure 5: Validation of RSPO2/3 rearrangements in oesophageal and biliary tract cancer cells.*** *(a) Sanger sequencing across fusion breakpoint for EIF3E-RSPO2 in ESO51 (an esophageal cancer cell line) and PTPRK-RSPO3 in EGI1 (a*

*biliary tract cancer cell line). (b) Fiber-FISH confirms presence of the EIF3E-RSPO2 fusion in ESO51. (c) FISH shows the presence of the PTPRK-RSPO3 fusion in EGI-1(arrows). (d) EGI1 and ESO51 are outliers for RSPO2 and RSPO3 gene expression in biliary tract and esophageal cancer cell lines, respectively. We identified an oesophageal cancer sample with high RSPO3 expression and a PTPRK-RSPO3 fusion. Z-score RNAseq values for TCGA samples were downloaded from cBioPortal. (e) Cell viability assay on EGI1 and ESO51 cells treated with the PORCN inhibitor WNT-C59 for 7 days. Data are expressed as average ± SD of three technical replicates from one representative experiment. SNU1411 is a positive-control colorectal cancer cell line with a known PTPRK-RSPO3 fusion. HCT116 is a negative-control colorectal cancer cell line. (f) The RSPO-fusions in ESO51 and EGI-1 show no differential essentiality to mapping versus non-mapping guides in the CRISPR drop-out screen.*

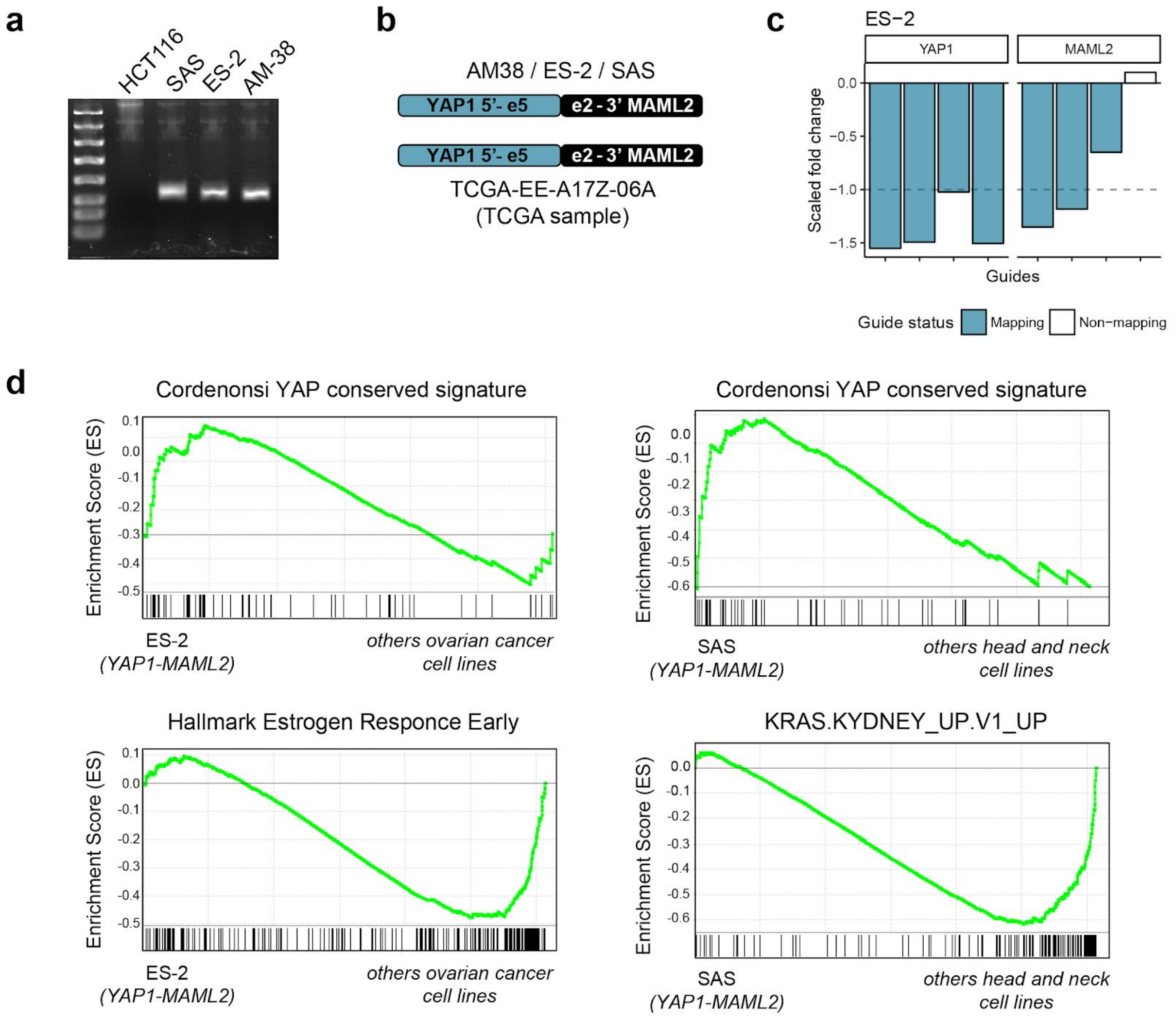


***Supplementary Figure 6: YAP1-MAML2 is a recurrent fusion required for cell fitness.*** *(a) PCR validation of the YAP1-MAML2 fusion in SAS, ES-2 and AM-38. HCT116 is a fusion-negative control. (b) The breakpoint of YAP1-MAML2 is preserved in all three cell lines, as well as in a patient sample identified in TCGA[2]. (c) Guides targeting the fused genes are differentially depleted depending on the fusion-mapping status. This holds true across multiple data resources, shown in two independent datasets for ES-2. Broad Institute data shown here and data from Sanger CRISPR screen shown in Figure 6. (d) Gene set enrichment analysis (GSEA) of YAP1 gene signature in YAP1-MAML2 positive ovarian (upper panel, left) and head and neck (upper panel, right) cancer cells vs fusion negative cell lines of the same tissue type. In the fusion-positive ovarian cancer cell lines ES-2, the gene*

*signature for early estrogen response is downregulated, compared to other ovarian cancer cell lines (left). In the fusion-positive head and neck carcinoma cell line SAS, the KRAS signature is downregulated, with respect to other head and neck carcinoma cell lines (right). The estrogen and the KRAS gene signatures are typical transcriptional hallmarks of the respective cancer types.*
